# Nanobubble based sonobiopsy reveals circulating protein signatures of BBB opening and glioblastoma

**DOI:** 10.64898/2025.12.04.692317

**Authors:** Roni Gattegno, Divsha Sher, Or Zohar, Keren Moskov, Dinorah Friedmann-Morvinski, Tali Ilovitsh

## Abstract

Focused ultrasound (FUS) and microbubbles can transiently increase blood brain barrier (BBB) permeability, yet verification of BBB opening (BBBO) relies mainly on contrast-enhanced MRI, offering limited insight into the molecular consequences of barrier modulation. Sonobiopsy, which uses FUS induced BBBO to release brain derived molecules into the bloodstream, provides a molecular readout from blood samples. Nanobubbles (NBs) are smaller agents that circulate more effectively in the brain microvasculature and have shown enhanced BBBO in capillaries. Here, NB-mediated proteomic sonobiopsy is used to improve biomarker efflux and increase molecular sensitivity, with the goal of defining the molecular signature produced by BBBO in healthy and glioblastoma (GBM) models, and distinguishing biomarkers associated with tumor pathology from BBBO biomarkers. Plasma collected before and after NB-mediated FUS underwent data-independent acquisition-based mass spectrometry, revealing post-BBBO changes in healthy and tumor bearing mice. In healthy cohorts, 77 proteins were reproducibly altered after BBBO. Six proteins (Dpysl3, Myl1, Mybpc1, Vsig4, Krt33a, Krtap6-5) were detectable only after BBBO in healthy mice. In 005 glioma bearing mice, the BBBO signature identified in healthy animals was preserved, and comparison with matched shams isolated tumor specific effects. Three proteins, Itih4, Lrg1, and Hp, rose significantly after BBBO only in GBM and reached higher post treatment levels than in shams, nominating candidate GBM-associated markers accessible via blood. These findings establish NB-mediated proteomic sonobiopsy as a promising method for BBBO verification and for detecting GBM associated protein signals, supporting the development of scalable BBBO confirmation and protein-based diagnostics in neuro oncology.

**Graphical Abstract:** 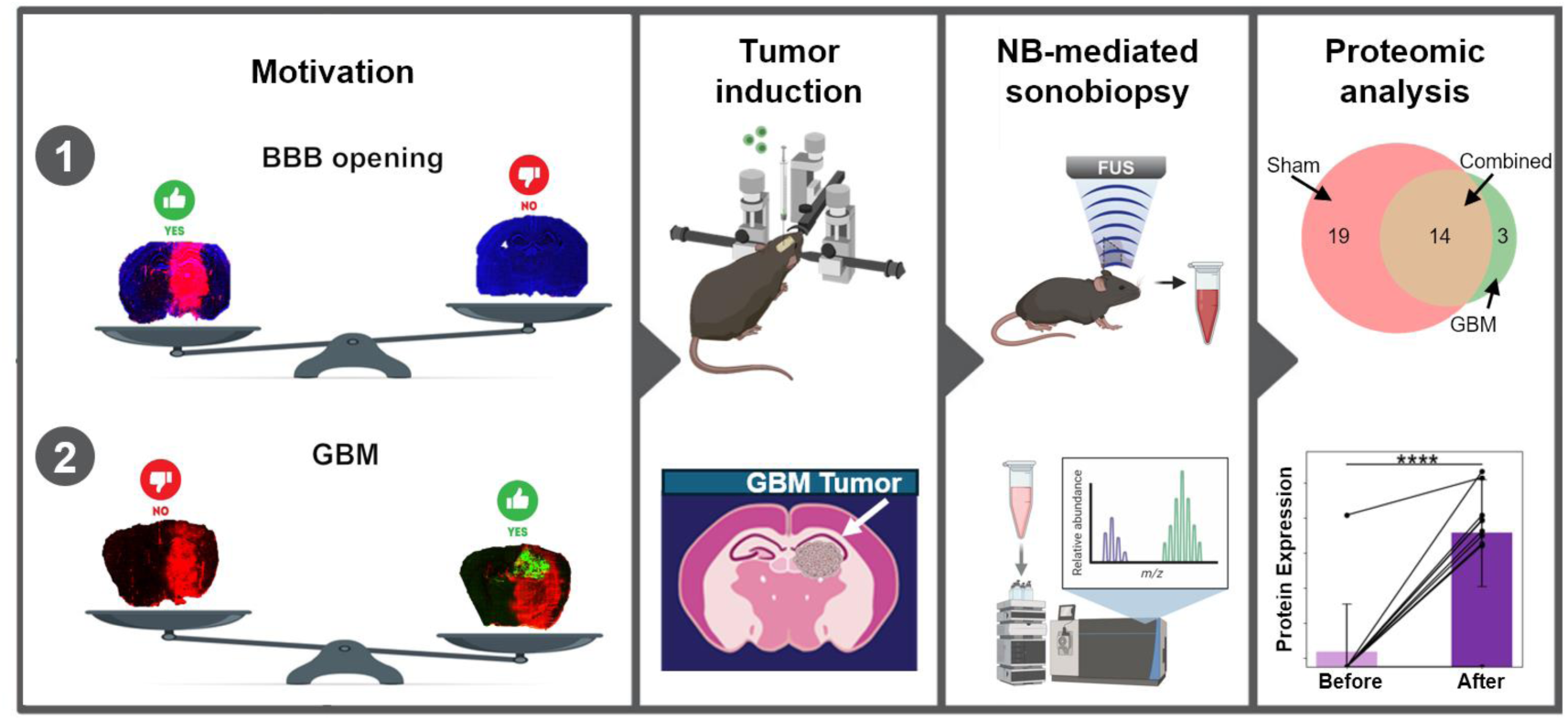

## Introduction

Focused ultrasound (FUS)-mediated blood brain barrier opening (BBBO) has emerged as a promising noninvasive technique to transiently increase the permeability of the blood brain barrier (BBB), facilitating the targeted delivery of systemically administered therapeutics to the central nervous system^1–5^. To ensure the clinical utility and reproducibility of FUS-based interventions, it is essential to characterize the physiological effects induced by BBBO and to develop reliable methods to confirm that BBB disruption has occurred in each treatment session. In this context, sonobiopsy is a molecular technique that leverages focused ultrasound (FUS) mediated BBBO to noninvasively sample brain-derived biomarkers from the bloodstream. By temporarily disrupting the BBB, molecules such as DNA, RNA, proteins, and extracellular vesicles, normally confined to the brain, can pass into circulation and be captured through a simple blood draw^6,7^. While originally developed with a focus on improving diagnostic sensitivity for brain pathologies^8–10^, sonobiopsy also holds promise as a general platform for evaluating BBB permeability through molecular signatures, a direction that has received limited attention to date.

While early applications of sonobiopsy have primarily centered on nucleic acid-based biomarkers such as circulating tumor DNA, cell-free DNA, or RNA transcripts, proteins may offer several distinct advantages. As the direct functional products of gene expression, proteins can provide real-time insight into the physiological state of the brain, including responses to mechanical stress or therapeutic interventions^11^. They are often more stable than fragmented nucleic acids, making them well-suited for detection in peripheral blood^12^. Moreover, by evaluating a broad panel of proteins rather than focusing on individual markers, it is possible to capture dynamic and heterogeneous changes that may not be apparent at the transcript level, ultimately offering a more comprehensive and accurate picture of brain state^13^. Although a few studies have begun to explore protein-based approaches, such as glial fibrillary acidic protein, myelin basic protein^6^, sonobiopsy remains largely focused on nucleic acids, and the full potential of proteomic profiling as both a diagnostic tool and a method for verifying BBBO has yet to be fully realized.

Given the importance of reliably releasing brain derived proteins into the bloodstream, the physical characteristics of the ultrasound contrast agents used during BBBO play a crucial role. To date, sonobiopsy studies have relied on BBBO using microbubbles (MBs)^6,8,9,14,15^. When FUS is applied to MBs with a typical diameter of 1-10 μm, it causes them to expand and contract repeatedly^16–18^. As this process occurs within blood vessels in the brain, the BBB can open safely and transiently to facilitate biomarkers release into the bloodstream. More recently, we have shown that nanobubbles (NBs) can enhance BBBO in capillaries^19–21^. NBs, which are significantly smaller, with diameters around 200 nanometers, can maneuver more easily within smaller vessels, leading to a more uniform BBBO in large and small blood vessels^20,21^. Recently, NB-mediated BBBO was successfully employed for the therapeutic delivery of genes and lipid nanoparticles^22,23^. To date, NB-based sonobiopsy has not been investigated, especially in combination with proteomic blood analysis. Because NBs improve BBBO in capillaries, we propose that this enhanced permeability could also increase the release of brain derived biomarkers into circulation. Our goal is to establish NB-based sonobiopsy and address two questions. First, we will test whether NB-based sonobiopsy can identify BBBO through a molecular readout in both healthy and tumor bearing mice. Second, after defining signatures unique to BBBO, we will determine which protein signals are associated specifically with the pathological condition.

While NB-based sonobiopsy offers a new molecular approach, we must evaluate it relative to existing validation techniques. Currently, most methods used to verify successful BBBO provide physical confirmation of barrier disruption, yet they differ significantly in clinical applicability and information depth. In preclinical studies, extravasation of dyes such as Evans Blue (EB) or fluorescent tracers is commonly used to confirm BBB permeability^17,18^. Although these approaches provide high-resolution spatial information, they require terminal tissue collection and are not applicable in clinical settings. In vivo imaging, particularly contrast-enhanced MRI, is widely used as a noninvasive alternative^24–28^. MRI enables real-time visualization of gadolinium leakage across the BBB and can be used to estimate the volume and duration of barrier opening. However, MRI is costly and not always available in all clinical or research environments. Moreover, in scenarios involving repeated FUS treatments with fixed targeting, such as in patients with implanted neuronavigation or robotic positioning systems, the need for MRI-guided targeting is reduced, yet there remains a critical need to confirm that the BBB has opened^29^. Ultrasound-based techniques, such as passive cavitation detection or power doppler imaging, offer a more accessible and lower-cost alternative for monitoring BBBO in real time^30–32^. However, similar to MRI, ultrasound-based techniques provide biophysical rather than molecular readouts and thus do not capture the downstream biological effects or mechanistic consequences of BBBO. In this study, we address this limitation by investigating NB-based sonobiopsy as a molecular method to verify BBBO directly from a blood sample.

In addition to serving as a tool for BBBO validation, protein signatures captured through sonobiopsy may also support diagnostic applications. Glioblastoma multiforme (GBM), the most aggressive primary brain tumor, remains difficult to diagnose and monitor due to the limitations of both invasive tissue biopsy and the molecular ambiguity of standard imaging^33,34^. While liquid biopsy offers a noninvasive alternative, its diagnostic sensitivity is limited by the low abundance of tumor-derived biomarkers in the bloodstream^35–37^. By enhancing the release of brain-specific proteins into circulation, sonobiopsy may facilitate the identification of GBM-associated protein biomarkers, thereby complementing existing nucleic acid-based approaches and broadening the scope of molecular diagnostics in neuro-oncology^15,38,39^.

To evaluate this, we utilized a mouse model of GBM alongside healthy controls, and we performed plasma proteomic analysis before and after BBBO to identify proteins released into circulation post BBBO. This comparative design allowed us to distinguish proteins associated with the BBBO procedure itself from those linked specifically to tumor pathology. By characterizing these protein signatures, our goal is to assess the potential of proteomics as a means of validating BBBO and to explore its utility in identifying candidate biomarkers for GBM.

## Results

### Global proteomic changes following FUS-mediated BBBO

To characterize the molecular consequences of BBBO in healthy mice, FUS-mediated BBBO was induced using intravenously administered NBs, and blood serum samples were collected a day before and 30 minutes after the procedure. Serum samples were subjected to mass spectrometry-based proteomic profiling, resulting in the detection of 1,754 proteins. Differential analysis revealed widespread changes in protein abundance (Fig. 1A), with 10 proteins showing both high statistical significance (p < 0.001) and large fold changes (FC, log₂FC > 3.5). To gain functional insight into the biological processes affected by BBBO, proteomaps were generated based on the fold change between pre- and post-treatment conditions. The high-level proteomap (Fig. 1B) revealed prominent enrichment in pathways related to central carbon metabolism, biosynthesis, protein folding and degradation, cytoskeletal remodeling, and vesicular transport, suggesting a systemic physiological response. A more detailed sub-pathway view (Fig. 1C) showed specific upregulation of proteins involved in glycolysis, tight junction modulation, cytoskeletal regulation, cell cycle progression, and exosome formation. This indicates that BBBO triggers acute biological responses involving metabolic activation, cellular restructuring, and intercellular signaling.

**Figure 1.**
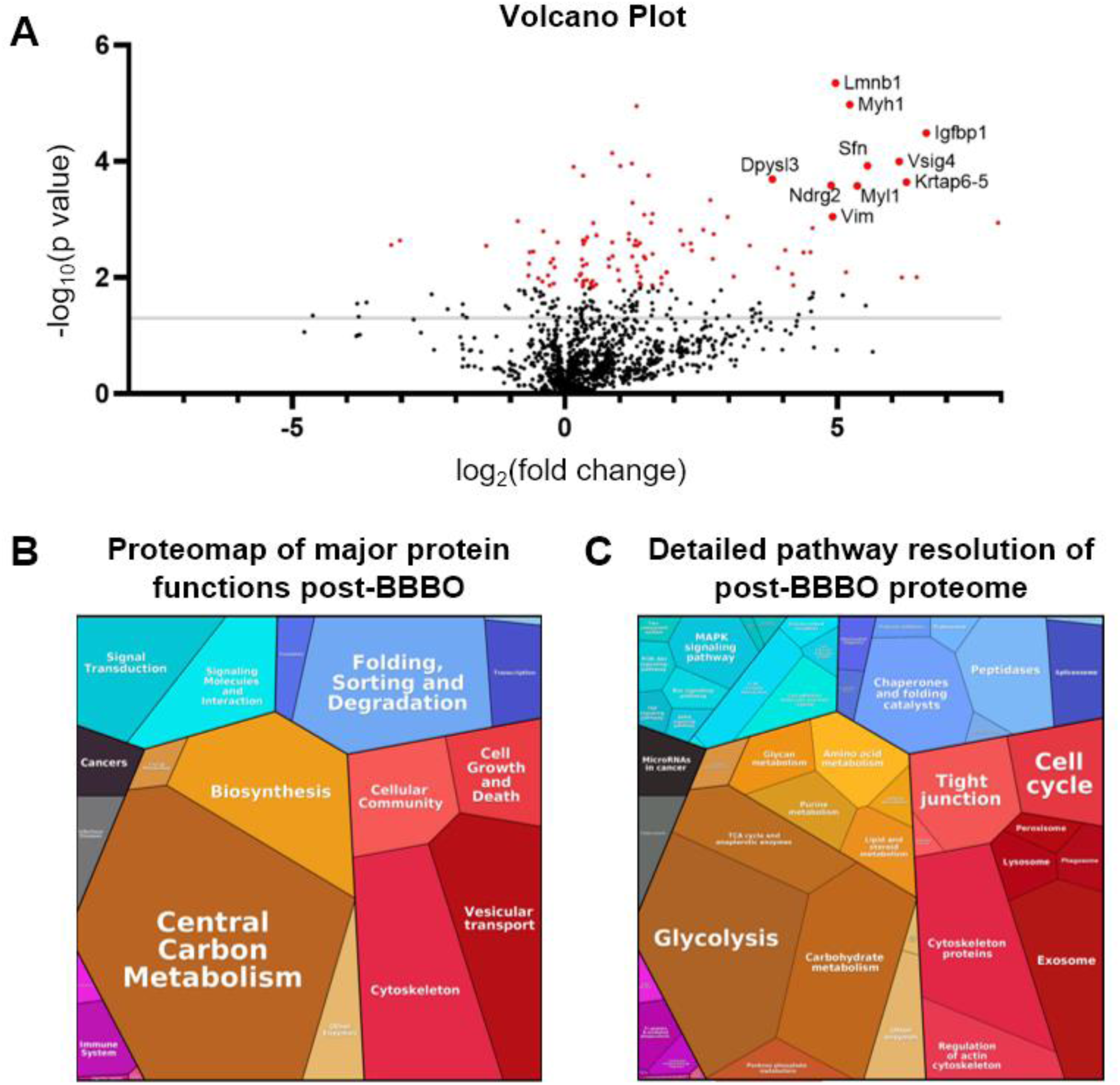
Proteomics overview in healthy mice following BBB opening. (A) Volcano plot showing differentially expressed proteins in healthy mice after BBBO. The grey horizontal line represents a −log₁₀(p-value) of 1.3 (corresponding to p = 0.05). Proteins with p-values < 0.01 are highlighted in red. Gene names are annotated for the 10 most significant proteins with a fold change greater than 3.5. (B) Proteomap illustrating high-level functional categories of proteins elevated post-BBBO. High-level biological processes are visualized based on fold change between post- and pre-BBBO conditions. (C) Proteomap illustrating sub-pathway-level functional enrichment among proteins upregulated after BBBO, based on fold change.

### Identification of robust protein markers associated with BBBO

To identify a consistent set of BBBO-associated proteins, two independent proteomic experiments in healthy mice were performed, five mice per group. In each experiment, pre-and post-BBBO serum samples were compared using paired *t*-tests. Proteins with a p-value < 0.05 and fold change > 1.5 were considered differentially expressed. Applying these selection criteria across both experiments yielded a subset of 77 proteins (Fig. 2A). To better understand the distribution of fold changes within this reproducible protein set, the proteins were divided into two groups: those exhibiting large expression changes (log₂FC > 2; Fig. 2B) and those with moderate changes (1.5 < log₂FC < 2; Fig. 2C). Among the highly regulated proteins (log₂FC > 2), 24 were common to both experiments and are listed in Table 1. Notably, six of these, Dpysl3, Myl1, Mybpc1, Vsig4, Krt33a and Krtap6-5, were detected exclusively in the post-BBBO samples across both experiments. The expression profiles for the second experiment, with proteins shown in the same order, are provided in Supplementary Fig. 1. Corresponding quantitative data for both experiments are available in supplementary tables 1 and 2, respectively.

**Figure 2.**
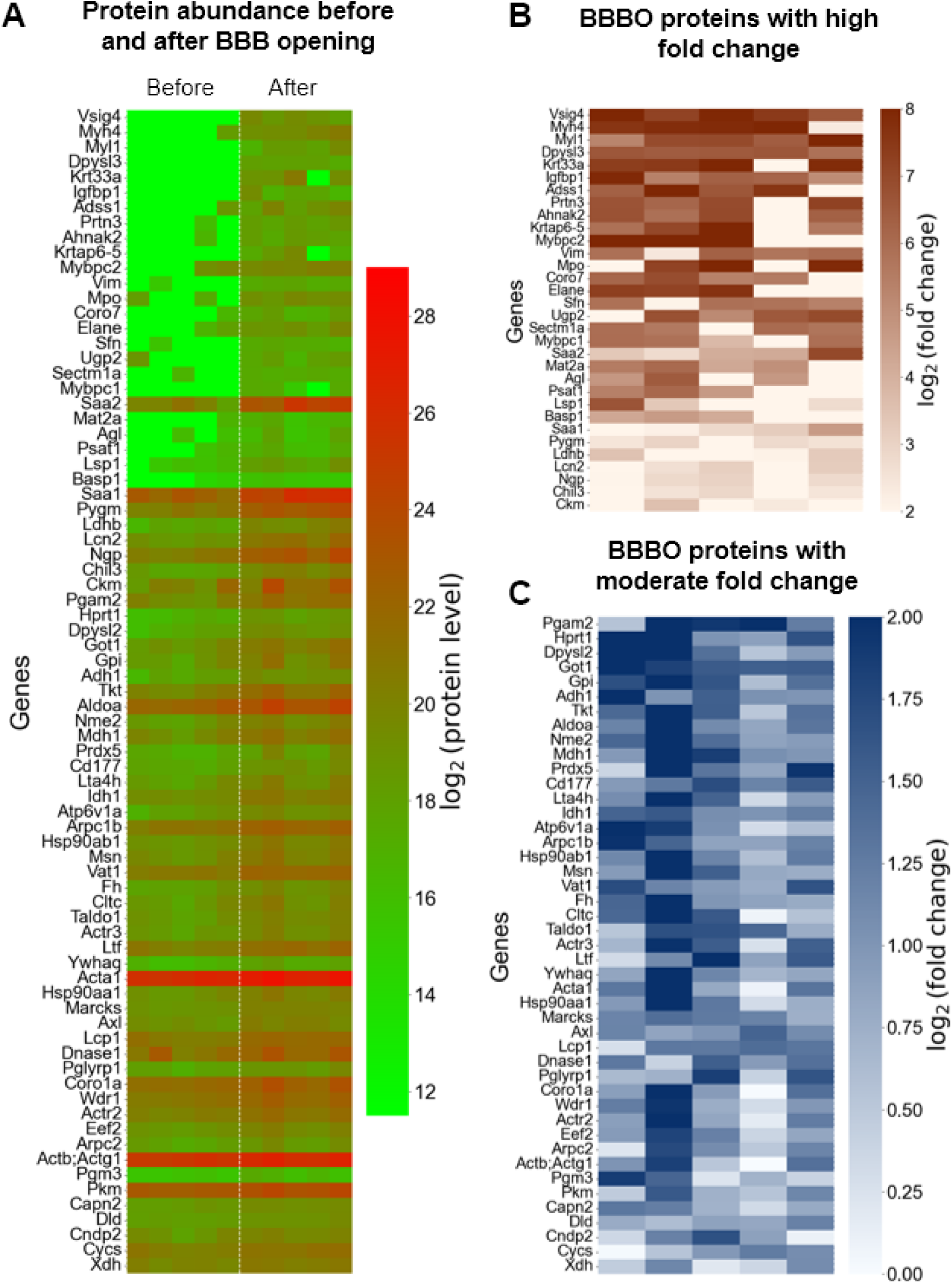
Proteomic markers associated with BBBO in healthy mice. (A) Heatmap showing log₂ transformed protein abundance for 77 proteins consistently altered following FUS-mediated BBBO in healthy mice. Differential expression was assessed independently in two experiments using paired t-tests comparing pre- and post-BBBO samples. Proteins with a p-value < 0.05 and fold change > 1.5 in both experiments were considered significant. The 77 overlapping proteins are presented here using data from the first experiment, and the corresponding values are provided in supplementary table 1. (B) Subset of BBBO-associated proteins exhibiting large fold changes (log₂ fold change ≥ 2). (C) Subset of BBBO-associated proteins exhibiting moderate fold changes (log₂ fold change < 2).

**Table 1.** Differential expressions of 24 proteins identified as significant in two independent sonobiopsy experiments in healthy mice. Numbers represent mean ± standard deviation. .

| Genes | log<sub>2</sub>FC 1<sup>st</sup> Experiment | log<sub>2</sub>FC 2<sup>nd</sup> Experiment |
| --- | --- | --- |
| Vsig4 | 7.46±0.64 | 7.64±0.22 |
| Myh4 | 6.82±2.54 | 8.00±0.45 |
| Myl1 | 6.73±1.00 | 6.99±0.20 |
| Dpysl3 | 6.40±0.32 | 4.76±0.42 |
| Krt33a | 6.36±3.64 | 8.21±0.41 |
| Igfbp1 | 6.17±1.12 | 5.11±2.27 |
| Adss1 | 6.04±2.82 | 4.69±2.76 |
| Krtap6-5 | 5.33±3.16 | 7.89±0.91 |
| Mybpc2 | 5.29±4.02 | 6.48±4.02 |
| Vim | 5.26±1.61 | 4.98±0.50 |
| Elane | 5.12±3.09 | 3.84±2.26 |
| Sfn | 4.99±1.64 | 2.81±2.06 |
| Ugp2 | 4.84±3.45 | 4.13±1.94 |
| Sectm1a | 4.79±2.34 | 5.10±2.87 |
| Mybpc1 | 4.28±2.57 | 2.76±1.77 |
| Saa2 | 4.28±1.64 | 4.96±1.88 |
| Mat2a | 4.17±2.36 | 2.09±1.39 |
| Agl | 3.48±2.76 | 5.17±0.84 |
| Psat1 | 3.42±2.74 | 2.49±1.72 |
| Saa1 | 2.88±1.16 | 3.94±2.19 |
| Pygm | 2.49±0.53 | 4.01±0.53 |
| Lcn2 | 2.39±0.86 | 2.25±0.37 |
| Ngp | 2.37±0.76 | 2.04±0.42 |
| Ckm | 2.04±0.96 | 2.27±0.71 |

### Cross-experimental reproducibility of BBBO-associated protein signatures

We next evaluated whether the identified protein signature could distinguish samples collected before and after BBBO across independent experiments. Principal component analysis (PCA) showed clear separation between pre- and post-BBBO samples from both datasets when projected into a common feature space (Fig. 3A). To assess whether a minimal protein panel could generalize across experiments, the 10 proteins with the largest FC among those significantly altered (p < 0.05) in the first experiment were selected. These proteins were selected solely based on their performance in the first dataset and were not required to overlap with the 77 shared proteins. A PCA model trained on this reduced feature set successfully preserved group separation when applied to the second experiment (Fig. 3B), suggesting that a compact biomarker panel may be sufficient to classify BBB status in independent cohorts.

**Figure 3.**
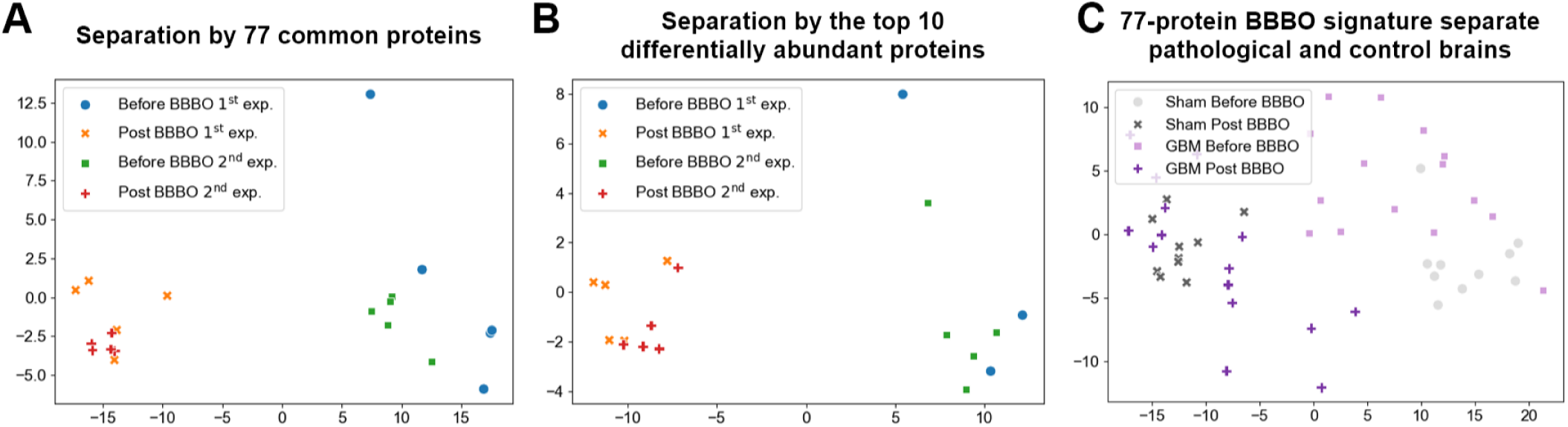
Proteomic sonobiopsy signatures consistently separate samples before and after NB-mediated BBBO. (A) PCA of the 77 proteins identified as significantly elevated following BBBO in two independent experiments. Samples from both experiments are projected using the same 77-protein feature set, demonstrating clear separation between pre- and post-BBBO conditions. (B) Cross-experiment generalization of BBB-opening markers. The top 10 proteins with the largest fold changes in the first experiment (not limited to the 77 shared proteins) were used to fit a PCA model, which was then applied to the second experiment. Despite the reduced feature set and independent sample source, pre- and post-BBBO samples from the second experiment remain clearly separable. (C) Sham and 005 glioma-bearing mice are projected using the same 77 BBBO-associated proteins identified in the healthy experiments, illustrating separation between pre- and post-BBBO samples and enabling comparison of BBBO responses in GBM versus sham conditions.

### Verification of BBBO in tumor-bearing mice using histological analysis

In order to test sonobiopsy in a pathological condition, we first confirmed BBBO using NBs in our disease model. The FUS protocol was applied to 005 glioma-bearing mice while sham group consisted of tumor-bearing mice without FUS treatment. Hematoxylin and eosin (H&E) staining confirmed the presence of tumor tissue in both control and FUS-treated groups (Fig. 4A-B). Since glioma vasculature can exhibit baseline leakiness, we systemically administered EB dye to evaluate whether FUS further enhanced vascular permeability beyond that associated with the tumor itself. Modest dye accumulation was observed in untreated controls, consistent with the leaky nature of glioma vasculature, whereas robust extravasation was evident in FUS-treated animals (Fig. 4C-D).

**Figure 4.**
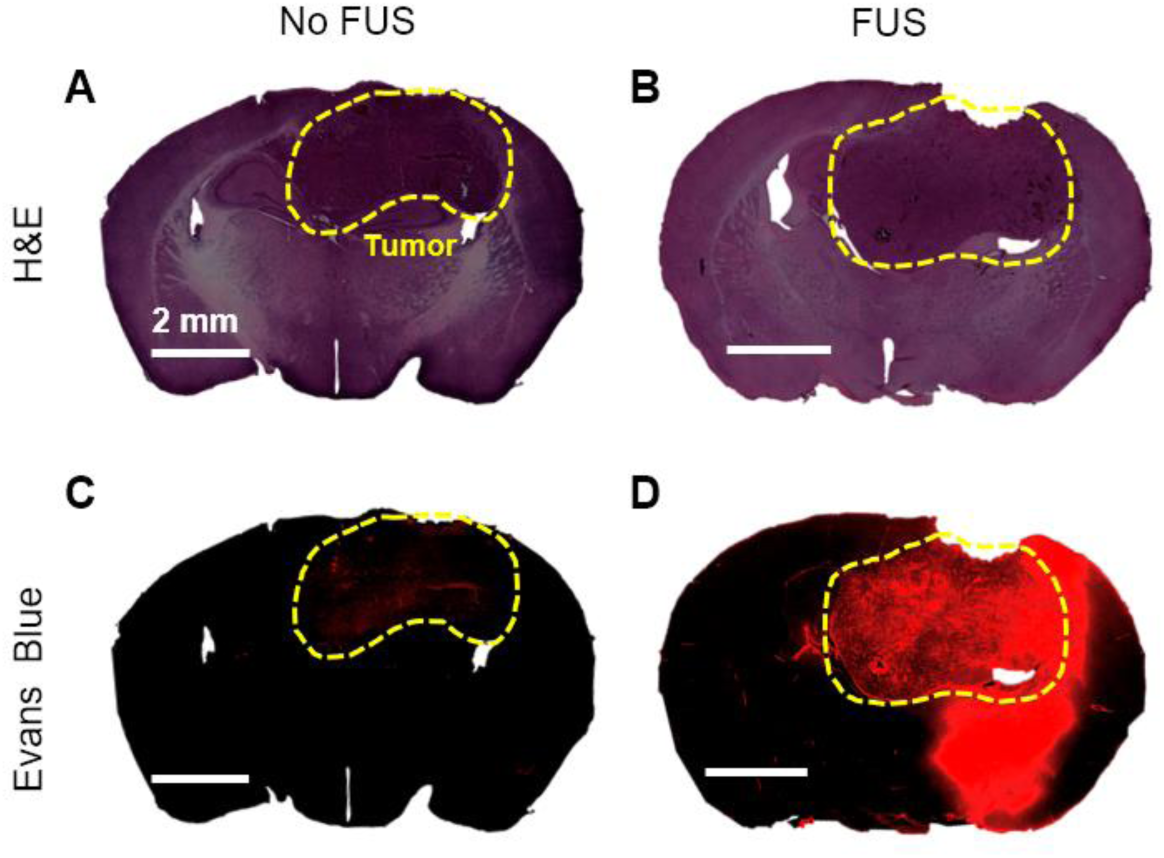
BBBO in 005 glioma-bearing mice. Representative coronal brain slices from 005 glioma-bearing mice. Hematoxylin and eosin (H&E) stained sections from (A) control (no FUS) and (B) NB + FUS groups. (C, D) Corresponding EB fluorescence images confirming BBB opening only in FUS-treated mice. Images were acquired with a 20× objective lens. Scale bar: 2 mm.

### Validation of candidate protein markers in sham and GBM models

Using the 77-protein BBBO signature derived from healthy mice, principal component analysis also separated pre- and post-BBBO samples in both sham and 005 GBM-bearing cohorts (Fig. 3C), indicating that the global opening signature generalizes to the tumor model. Among the upregulated proteins, we previously identified six candidates (Dpysl3, Myl1, Mybpc1, Vsig4, Krt33a, and Krtap6-5) that consistently appeared only after BBBO in healthy mice (Table 1). To validate these candidates also in the GBM model, we quantified their serum expression before and after FUS-mediated BBBO in both sham- and GBM-bearing animals (Fig. 5). In sham-treated animals, all six proteins showed significant induction following BBBO: Dpysl3, Myl1, Vsig4, Krt33a, Krtap6-5 (p < 0.0001) and Mybpc1 (p < 0.01). In the GBM cohort, these proteins demonstrated a similar trend, with significant increases observed for Myl1, Krt33a, Krtap6-5 (p < 0.0001), Vsig4 (p < 0.001), Dpysl3 and Mybpc1 (p < 0.01). Although variability was greater in tumor-bearing animals, these proteins had similar trends compared to sham. The repeated upregulation supports their robustness as markers of BBBO across both healthy and pathological contexts and their potential as indicators of BBBO.

**Figure 5.**
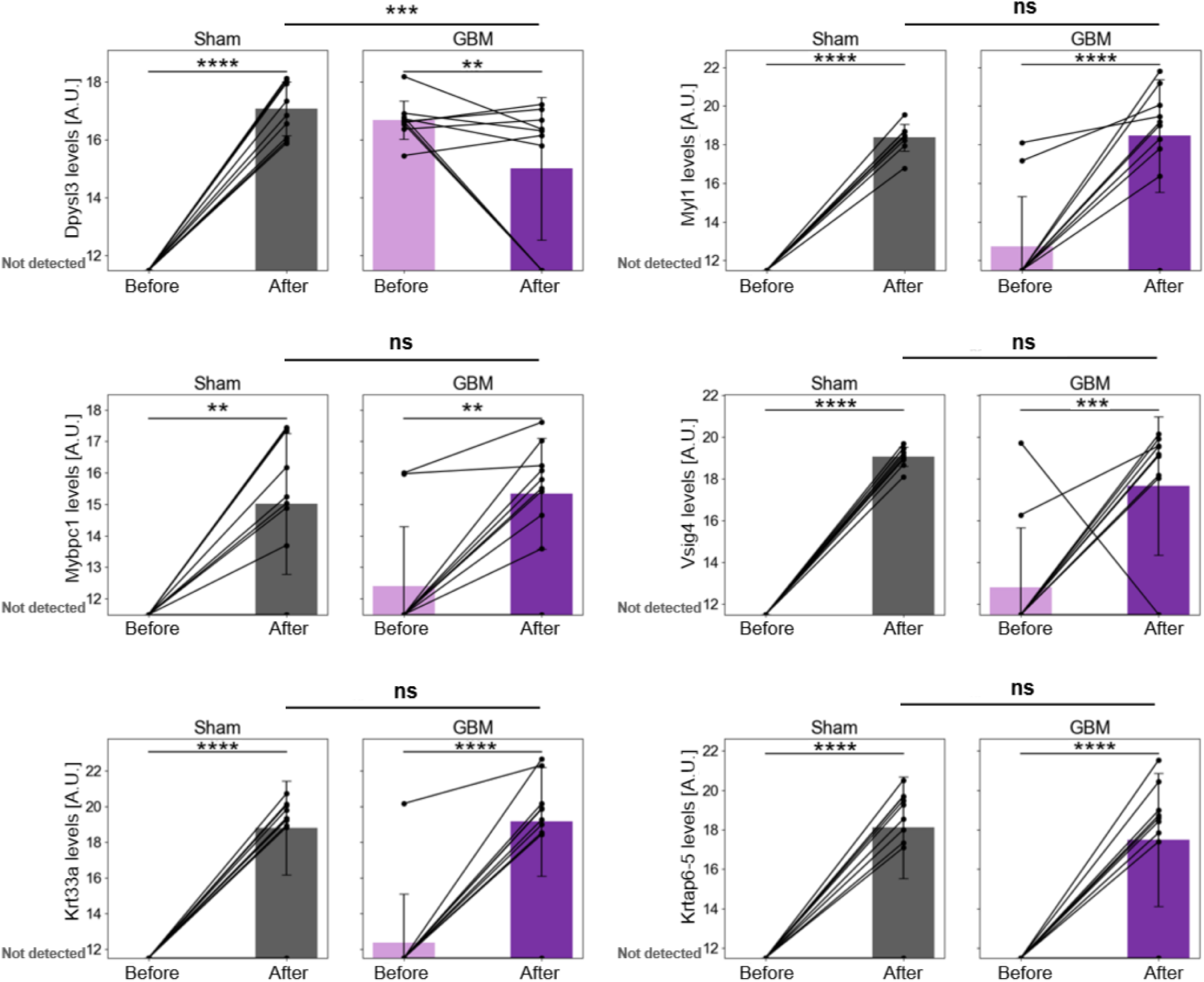
Proteins detected only after BBBO in sham and GBM cohorts. Line plots show changes in expression levels of six proteins that were detectable only after BBBO in the sham-treated group; (A) Dpysl3, (B) Myl1, (C) Mybpc1, (D) Vsig4, (E) Krt33a, and (F) Krtap6-5. Left subplots represent sham-treated animals, and right subplots represent GBM-treated animals. Bars represent group means ± SD; light bars indicate *Before*, dark bars indicate *After*. Each dot represents an individual animal, with lines connecting repeated measures. Statistical significance was assessed using two-way ANOVA with Tukey correction for multiple comparisons, with significance denoted as: p < 0.05 (*), p < 0.01 (**), p < 0.001 (***), p < 0.0001 (****), and not significant (ns).

### Identification of GBM-specific candidate biomarkers

To isolate tumor-context effects from procedure-related effects, proteins whose post-BBBO changes were specific to GBM were searched. For each condition (sham, GBM), we first applied a consistency filter across the two independent experiments: proteins were retained only if they were either significant in both experiments (p < 0.05 and FC>1.5, post vs. pre) or non-significant in both. Among these, 14 proteins were shared by both conditions, 19 were unique to sham, and 3 were unique to GBM (Fig. 6A). The three GBM-unique proteins, inter-α-trypsin inhibitor heavy chain 4 (Itih4), Leucine-rich α-2 glycoprotein 1 (Lrg1), and Haptoglobin (Hp), showed significant increases after BBBO in GBM-bearing mice (Lrg1: p < 0.001; Itih4 and Hp: p < 0.01), whereas no significant changes were observed in sham controls. Furthermore, post-treatment levels of Lrg1 and Hp were significantly higher in GBM compared to sham (p < 0.05 and p < 0.01, respectively) (Fig. 6B-D). Itih4, Lrg1, and Hp may reflect GBM-associated protein responses to FUS-mediated BBBO and could serve as candidate biomarkers for glioma-related modulation.

**Figure 6.**
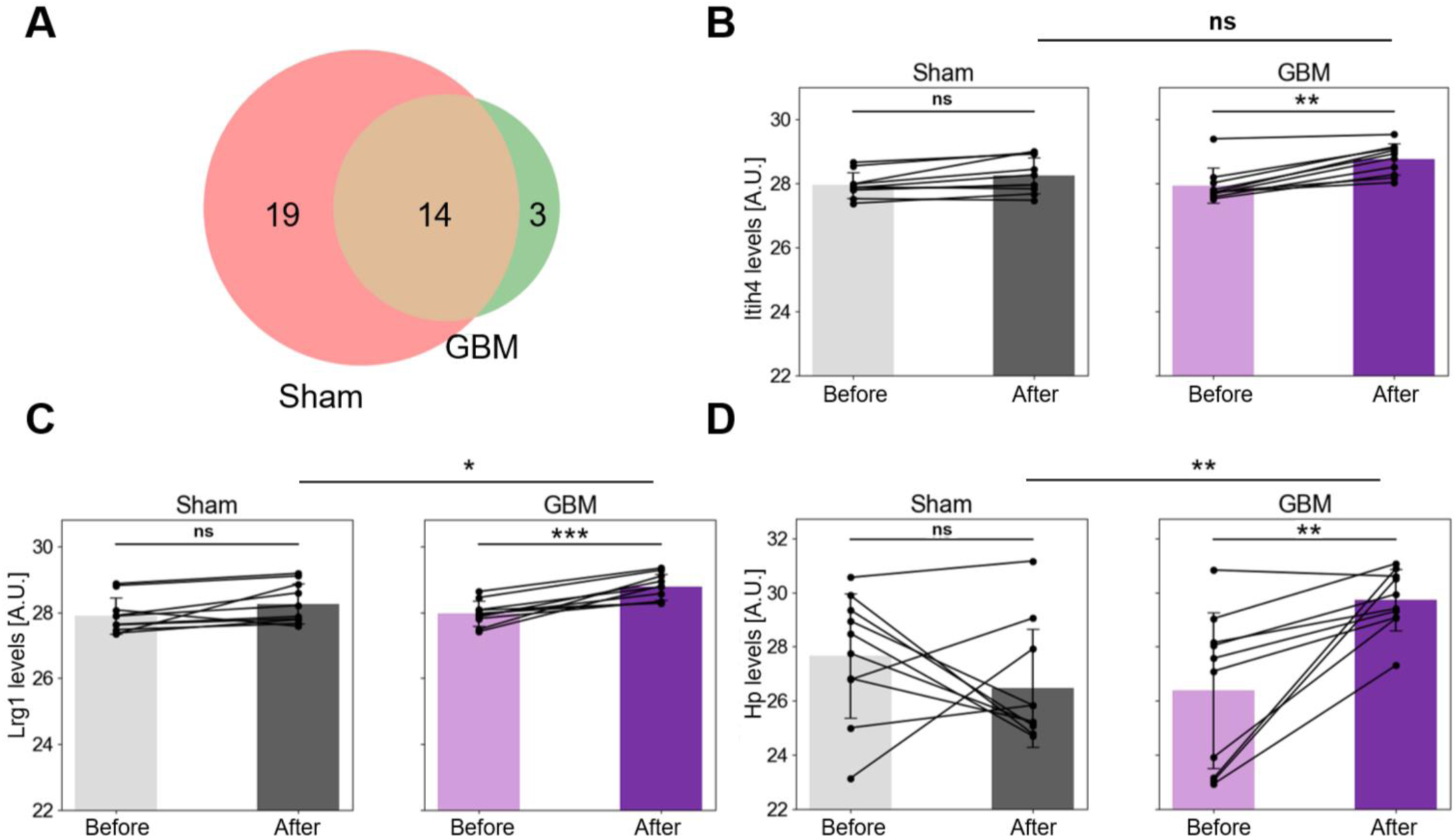
Post-BBBO proteins significantly elevated exclusively in GBM. (A) Venn diagram showing the overlap between detected proteins in Sham and GBM groups. Protein levels before and after treatment for (B) Itih4, (C) Lrg1, and (D) Hp. Left subplots represent sham-treated animals, and right subplots represent GBM-treated animals. Bars represent group means ± SD; light bars indicate *Before*, dark bars indicate *After*. Each dot represents an individual animal, with lines connecting repeated measures. Statistical significance was assessed using two-way ANOVA with Tukey correction for multiple comparisons, with significance denoted as: p < 0.05 (*), p < 0.01 (**), p < 0.001 (***), p < 0.0001 (****), and not significant (ns).

## Discussion

Here, we establish and evaluate NB-mediated proteomic sonobiopsy as a tool to detect blood-based biomarkers released after FUS-mediated BBBO. Using this framework, we address two questions: whether plasma proteins can provide a reproducible molecular signature of BBBO in healthy mice, and whether they can reveal GBM-associated signals when applied in a tumor model with matched shams. Across two independent experiments in healthy animals, 77 proteins changed consistently after BBBO, and a compact subset as few as 10 proteins alone was sufficient to classify samples before vs. after opening. The FUS parameters used here were effective for inducing BBBO also in tumor-bearing brains. The 005 glioma model was selected because it recapitulates key molecular and histopathological features of human mesenchymal GBM and retains an intact BBB at early stages^40,41^, providing a suitable platform to assess FUS-mediated BBBO. Consistent with this, Fig. 4C shows limited extravasation in non-sonicated tumors, indicating that the barrier remains relatively preserved without FUS. In tumor-bearing mice, we further observed post-BBBO protein changes that were preferentially associated with the GBM cohort compared with shams, indicating that tumor-related signals can be detected on top of the procedural effect of BBBO. These findings support the use of plasma proteomics both to verify BBBO and to reveal tumor-linked biology in the setting of sonobiopsy.

Previous studies employing MBs have reported protein- and transcript-level changes after FUS-mediated BBBO, most often involving acute inflammatory pathways^42–48^. However, work with NBs has largely focused on permeability, safety, and delivery efficiency rather than on downstream molecular responses^20–23,32,49,50^. Our findings extend this body of work by providing the first evidence that NB-FUS induces a reproducible proteomic signature detectable in circulation. In healthy mice, six proteins were detectable only after BBBO: Dpysl3, Myl1, Mybpc1, Vsig4, Krt33a, and Krtap6-5. Dpysl3 is a neuronal cytoskeleton and axon-guidance protein that is induced in activated microglia under inflammatory stimuli, consistent with a BBB-adjacent neuroinflammatory response surfaced by sonication^51^. Myl1 encodes a fast skeletal myosin light chain that is elevated in CSF after ischemic stroke and thus can appear in central compartments during vascular compromise^52,53^. Vsig4 is a complement-receptor family protein on tissue macrophages, including CNS border macrophages, and acts as a protective modulator of BBB integrity and neuroinflammation after intracerebral hemorrhage. Its post-BBBO detection is consistent with immune-vascular signaling during opening^54^. Although Mybpc1, Krt33a, and Krtap6-5 have not, to our knowledge, been linked directly to BBB permeability, their post-BBBO-only detection points to roles as permeability-sensitive markers that strengthen the composite circulating signature. Together, the consistent pre- vs post-BBBO separation across experiments and the identification of a compact set of markers suggests that circulating proteins can serve as reliable indicators of BBBO, with some markers reflecting neurovascular activation and others functioning as sensitive sentinels of increased permeability.

The ability to confirm BBBO at the molecular level has several potential applications. In preclinical studies, it could replace invasive histological dyes or tissue collection, enabling repeated measures in the same animal and thereby reducing within-subject variability and overall animal use. Clinically, molecular readouts can complement imaging: MRI will remain essential to localize the first sonication, but once targeting is established, serial molecular checks could reduce the need for follow-up MRIs, particularly in protocols with multiple sonications and fixed targeting platforms, or when access to MRI is limited^29^. Beyond verification, the temporal profile of these proteins may inform BBB status and closure kinetics. Similar markers may also report barrier permeability in other contexts, including electrically induced opening, chemical opening, and diseases with BBB compromise such as epilepsy^24,55–59^. It remains to be established whether the same proteins can serve as indicators not only of FUS-induced BBBO, but also of these additional conditions. Finally, because the released proteins reflect cellular and vascular responses to ultrasound, they may guide optimization of FUS parameters for safe and effective translation.

In addition to BBBO-related proteins, our design allowed us to isolate tumor-specific responses. Clinical studies of sonobiopsy have shown that BBBO enhances the detection of brain-derived molecules^15,38^, but comparisons with healthy controls are rarely feasible for ethical reasons. Here, by comparing GBM and sham mice, we distinguished tumor-associated changes from those attributable to the procedure alone. Three proteins, Itih4, Hp and Lrg1, were consistently induced only in GBM-bearing animals after BBBO, suggesting that they may reflect glioma-related biology rather than a generic barrier response. Itih4 is a liver-derived acute-phase glycoprotein and protease inhibitor^60,61^. While often interpreted as systemic, it has been detected in GBM in surgical aspirates, bulk tumor proteomics and increases under GnRH-agonist treatment in GBM cells, supporting microenvironmental relevance even if tumor-intrinsic production is uncertain^62–64^. Hp is the canonical hemoglobin-scavenger that drives CD163-mediated clearance and limits oxidative injury^65–68^. Consistent with the increase that sonobiopsy helps us detect, multiple studies report elevated serum Hp in GBM and several explicitly propose Hp as a GBM serum biomarker^69–72^. Notably, in our cohort, serum Hp rose after sonobiopsy in 9/10 GBM mice, whereas in shams it increased in only 4/10 and decreased in the remainder, indicating the post-BBBO rise is preferentially GBM-linked. Lrg1 is a secreted leucine-rich glycoprotein that reprograms TGF-β signaling to promote pathological angiogenesis^73,74^. Beyond being overexpressed in GBM^75–77^, Lrg1 levels are higher in GBM than in lower-grade gliomas such as astrocytoma and oligodendroglioma^78^, supporting its value alongside imaging to help characterize tumor biology. Together, these GBM-restricted post-BBBO increases show how a GBM-vs-sham design can distinguish tumor effects from procedure effects and reveal a concise serum-accessible signature. Scaling this approach could refine GBM-specific signature and may even uncover additional, under-recognized biomarkers for GBM.

Discovering novel proteins associated with GBM through NB-mediated sonobiopsy has several important implications. Early diagnosis and longitudinal monitoring remain challenging in neuro-oncology because conventional imaging lacks molecular specificity and circulating tumor DNA is often scarce in blood^33–35,37^. Protein biomarkers identified through this platform could improve sensitivity, complement nucleic-acid assays, and broaden the diagnostic toolkit^36^. Beyond GBM, sonobiopsy has been shown feasible across other neurological contexts, enabling systematic discovery of circulating protein biomarkers that are otherwise sequestered by the BBB^9^.

However, several limitations still need to be taken into consideration. Discovery-scale proteomic analysis is costly and often sample-limited, so translation will require targeted assays in larger cohorts. Although NB-mediated BBBO is more spatially confined than MB-mediated opening^23^, enabling finer targeting and better regional precision, the treated field can still extend beyond the tumor. Greater focal selectivity is likely achievable by increasing FUS frequency in preclinical studies to tighten the focal geometry, and by using real-time feedback control to steer dosing toward tumor-selective opening. The kinetics of protein release after BBBO are not yet optimized as a single post-treatment time point was sampled. Finer time courses, including longer intervals between sonication and blood collection, may capture additional proteins which are required to monitor BBB closure, since informative markers should clear from blood faster than the barrier reseals. Our analysis emphasized proteins that increased after opening. A complementary analysis of down-regulated proteins could reveal additional biology. A further limitation concerns the distinction between BBBO markers and GBM-associated proteins. Prior BBBO indicators often behaved in a binary fashion (detected vs. undetected), whereas the GBM-linked proteins we identified were detectable at baseline and showed a preferential rise after opening in GBM but not in shams, indicating subtler, context-dependent shifts. This difference motivates systematic time-course studies across tumor stages to separate baseline leakage from procedure-evoked release. Beyond cost, proteomics has inherent constraints in sensitivity, dynamic range, variability introduced before measurement during collection, processing, and storage. Once candidates are nominated, independent validation with targeted methods such as ELISA, PRM/MRM mass spectrometry, and Western blot, and, where appropriate, qPCR in matched tissues will be important. To establish disease specificity, future work should assess whether the GBM-linked proteins generalize across or differentiate from other primary brain tumors.

Altogether, our findings establish proteomic sonobiopsy with NBs as a dual-purpose platform: it provides a reproducible molecular signature of BBBO itself and enables the discovery of tumor-associated proteins in GBM. With standardized protocols, targeted assay development, and prospective validation, proteomic verification can reduce dependence on costly imaging for BBBO confirmation and expand the molecular diagnostic toolkit for GBM by enabling discovery and monitoring of clinically relevant biomarkers.

## Methods

### 005 mouse GBM models

005 glioma stem cell (GSC) line was used to create the mouse GBM model. It was generated from a lentiviral HRasV12-induced tumor in a p53−/− knockout mouse^40^. For in vivo monitoring, a luciferase-expressing (bioluminescent) 005 variant was used^79^, which enabled IVIS bioluminescence imaging. The 005 cells were cultured in DMEM F12 media (Gibco, Massachusetts, USA) supplemented with Glutamax (1:100, Gibco, Massachusetts, USA), 100 units/mL penicillin, 50 mg/mL streptomycin, N2 supplement (1:100, Gibco), 2.5 μg/mL heparin (Sigma, Germany), 20 ng/mL FGF (Peprotech, New Jersey, USA), and 20 ng/mL EGF (Peprotech, New Jersey, USA). The cultures were maintained at 37 °C with 5% CO₂.

Mice were housed under pathogen-free conditions at the Tel Aviv University Specific Pathogen Free facility before the stereotactic injection. To induce tumors, 300,000 005 cells were stereotactically injected into the hippocampus (AP = -2, ML = -1.5, DV = -2.3) of each mouse using microsyringes (Hamilton, cat. n87925, Nevada, USA) mounted on a stereotaxic instrument (KOPF model 900, California, USA). Following the injection, mice were monitored daily for signs of illness. Tumor size was assessed one day prior to the sonobiopsy procedure using the IVIS Spectrum system (PerkinElmer, Massachusetts, USA).

### Study design

In this study, a total of 20 male C57BL/6 mice (11.8 ± 2.8 weeks old, 23.7 ± 2.3 g; Envigo, Jerusalem, Israel) were used. The mice were divided into two groups: GBM and sham. All animal experiments were conducted in accordance with the Guide for the Care and Use of Laboratory Animals and were approved by the Institutional Animal Care and Use Committee at Tel-Aviv University. Each experiment (two in total) included five mice per group (GBM and sham, Fig. 7). On day 0, GBM-bearing mice received stereotactic injections of 005 glioma cells into the hippocampus, while sham mice received vehicle injection of Phosphate Buffered Saline (PBS). On day 19, GBM group was scanned using an IVIS imaging system. On day 20, both groups underwent the sonobiopsy procedure: NB-mediated BBBO was performed, followed by blood collection and serum extraction for proteomic analysis as detailed below. After blood collection, EB was administered, followed by euthanasia and brain harvest.

**Figure 7.**
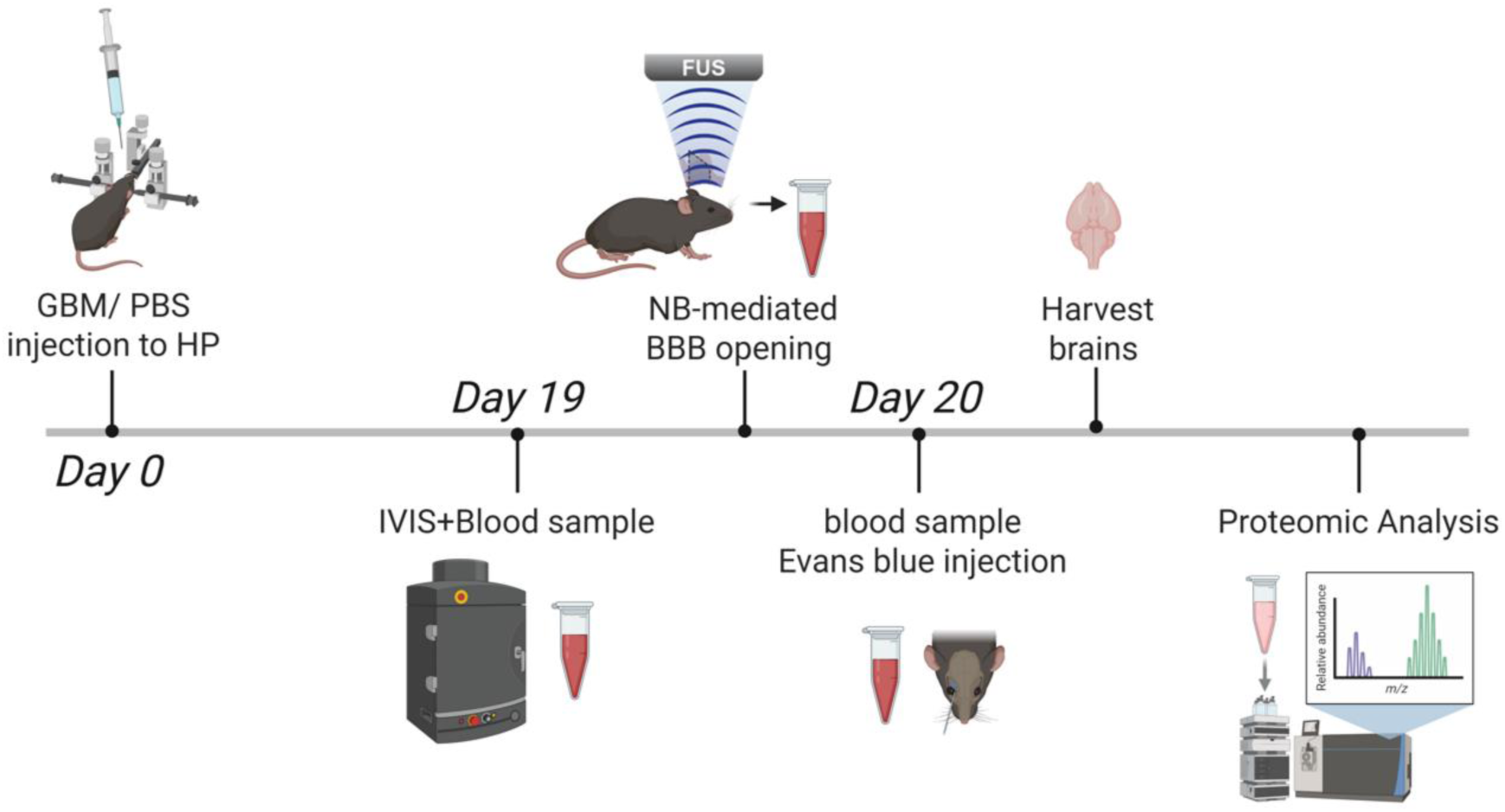
Study design. Overview of the sonobiopsy pipeline for noninvasive proteomic profiling following NB-mediated BBBO.

### Nanobubble preparation

NBs were prepared as described in earlier studies^20,80,81^. Briefly, NBs with an average diameter of 180 nm were fabricated with a phospholipid shell and an octafluoropropane (C_3_F_8_) gas core. A phospholipid solution (10 mg/mL) was prepared by dissolving 1,2-dibehenoyl-sn-glycero-3-phosphocholine (C22, Avanti Polar Lipids Inc.), 1,2-dipalmitoyl-sn-glycero-3-phosphate (DPPA, Corden Pharma), 1,2-dipalmitoyl-sn-glycero-3-phosphoethanolamine (DPPE, Corden Pharma), and 1,2-distearoyl-sn-glycero-3-phosphoethanolamine-N-[methoxy(polyethylene glycol)-2000] (DSPE-mPEG2000, Laysan Lipids) in propylene glycol (Sigma Aldrich).

The lipid solution was mixed with glycerol (Gly, AcrosOrganics) and phosphate buffer (pH 7.4) preheated to 80 °C, followed by sonication at room temperature for 10 minutes (Elmasonic P 60H, Elma Schmidbauer GmbH, Germany). The solution was transferred to a 2 mL vial (Milipore Sigma, Massachusetts, USA), sealed, purged with octafluoropropane (C_3_F_8_) gas (Apollo Scientific, Manchester, UK) and kept in 4 °C. Upon use, the NBs were activated by mechanical shaking (VialMix, Bristol-Myers Squibb Medical Imaging Inc., Massachusetts, USA) and separated through centrifugation (5810R centrifuge, Eppendorf AG, Hamburg, Germany), 50 rcf for 5 minutes. The lower phase was collected while the bubble cake was discarded, yielding a purified NB solution.

### Sonobiopsy Procedure

The sonobiopsy setup followed previous studies^20,82^, utilizing a spherical single-element FUS transducer (H115, Sonic Concepts, Bothell, WA, USA) submerged in a water tank. The transducer was driven by a transducer power output system (TPO-200, Sonic Concepts) connected through the manufacturer’s matching network and was operated at 850 kHz with a peak negative pressure of 210 kPa, producing a focal spot of 1.5 mm laterally and 6 mm axially to enable BBBO in the tumor region. The transducer was oriented upward at the bottom of the tank, and the acoustic path was coupled through an agarose phantom placed at the focus. The agarose (1.5% w/v in degassed water) was prepared by heating to dissolve, poured into a custom mold, and allowed to set before use. The mice, anesthetized with isoflurane, were aligned on the FUS setup using a 3D-printed head holder positioned on an agarose phantom, mounted on micrometer stages for fine positioning. Peak negative pressures at focus were calibrated with a needle hydrophone (NH0500, Precision Acoustics, UK). A retro-orbital injection of NBs (1 μL/g, diluted 1:1 in PBS) was administered, and FUS treatment commenced 30 seconds later to ensure optimal bubble distribution. The FUS protocol consisted of 1 ms bursts at 849 kHz with a PRF of 1 Hz (0.1% duty cycle) for 60 seconds. For serum extraction, blood was collected before EB injection into serum tubes, allowed to clot (20 minutes, room temperature), and centrifuged at 20,000 rcf for 10 min at 4°C in a benchtop centrifuge (5424R, Eppendorf, Hamburg, Germany). Serum was stored at −80 °C for subsequent proteomic analysis. For EB extravasation assays, a sterile 2% (w/v) EB solution in PBS was prepared and filtered (0.22 μm). EB was injected retro-orbitally (4 mL/kg) and allowed to circulate for 30 min. Mice were perfused 30 minutes after EB injection with PBS (0.01 M), followed by 4% paraformaldehyde. Brains were harvested, post-fixed at 4 °C, cryoprotected in 30% sucrose, embedded, and sectioned into 40 μm slices. Slices were stained with H&E to examine tissue integrity and damage. Fluorescence imaging was used to assess EB extravasation. Images were acquired using a motorized microscope (Revolution, Echo, San Diego, USA).

### Proteomic analysis

#### Proteolysis

The samples were precipitated in ice-cold TCA-Acetone (10% TCA and 90% Acetone) at a 1:4 ratio sample:TCA/acetone overnight at -20 °C and were washed with ice cold acetone. The protein pellets were dissolved in 8.5 M urea, 400mM ammonium bicarbonate, and 10mM DTT. Protein amount was estimated using Bradford readings. The samples were reduced (60 °C for 30 min), modified with 35.2 mM iodoacetamide in 100 mM ammonium bicarbonate (room temperature for 30 minutes in the dark) and digested in 1.5 M urea, 66 mM ammonium bicarbonate with modified trypsin (Promega), overnight at 37 °C in a 1:50 (M/M) enzyme-to-substrate ratio. An additional second trypsinization was done for 4 hours in a 1:100 (M/M) enzyme-to-substrate ratio. The tryptic peptides were acidified to 1% formic acid.

#### Mass spectrometry analysis

The peptides were analyzed by LC-MS/MS using an Exploris 480 mass spectrometer (Thermo Fisher, Massachusetts, USA) fitted with a capillary HPLC (EV-1000, Evosep One). The peptides were loaded onto a 15 cm, ID 150 µm, 1.9-micron Performance column EV1137 (Evosep). The peptides were eluted with the built-in Xcalibur 15 SPD (88min) method. Mass spectrometry was performed in a positive mode using repetitively full MS scan (m/z 380–985, resolution 120,000) followed by DIA scans (10 Da isolation windows with 1 m/z overlap, and resolution 30,000).

#### Data analysis

The mass spectrometry data was analyzed using the DIA-NN software version 1.8.1^83,84^ searching against the mus musculus proteome from the Uniprot database, with minimal peptide length set to 7, maximum number of missed cleavages set to 1, cysteine carbamidomethylation enabled as a fixed modification, and protein N-term acetylation enabled as a variable modification. Peptide- and protein-level false discovery rates (FDRs) were filtered to 1%. Statistical analysis of the identification and quantization results was done using Perseus 1.6.7 software^85^.

#### Statistical analysis

Statistical analysis was performed using Python and GraphPad Prism. Specific statistical tests are reported in the figure captions.

## Acknowledgements

This work was supported in part by the Israel Science Foundation under Grant 192/22, in part by an ERC StG under Grant 101041118 (NanoBubbleBrain), in part by the Israel Cancer Research Fund (grant number 1286686), in part by the Nicholas and Elizabeth Slezak Super Center for Cardiac Research and Biomedical Engineering at Tel Aviv University, and in part by the Adams Fellowship Program of the Israel Academy of Sciences and Humanities (R.G.). Proteomic analysis was performed by the Smoler Proteomics Center at the Technion Research and Development Foundation Ltd. Fig. 7 was created with BioRender.com (Publication License: https://BioRender.com/r5tmua2).

## Competing interests

The authors declare no competing interests.

## Contributions

R.G designed and performed the research, analyzed the data, and wrote the manuscript. D.S designed and performed the research. O.Z and K.M assisted with in-vivo experiments. D.F.M guided, advised and contributed to the manuscript. T.I. guided, advised, designed the research and contributed to the manuscript. All authors reviewed the manuscript.

## Declaration of generative AI use

During the preparation of this work the authors used ChatGPT (OpenAI) to assist with language editing and rephrasing certain sections of the manuscript. After using this tool, the authors reviewed and edited the content as needed and they take full responsibility for the content of the published article.

## Notes

### Competing Interest Statement

The authors have declared no competing interest.

