## Supplementary for "Nanobubble based sonobiopsy reveals circulating protein signatures of BBB opening and glioblastoma"

[1]The School of Biomedical Engineering, Tel Aviv University, Tel Aviv, Israel

[2]The Sagol School of Neuroscience, Tel Aviv University, Tel Aviv, Israel

[3]Department of Biochemistry and Molecular Biology, The George S. Wise Faculty of Life Sciences, Tel Aviv University, Tel Aviv, Israel

**This PDF file includes:**

**Supplementary Text**

**Fig. S1**

**Supplementary table 1**

**Supplementary table 2**

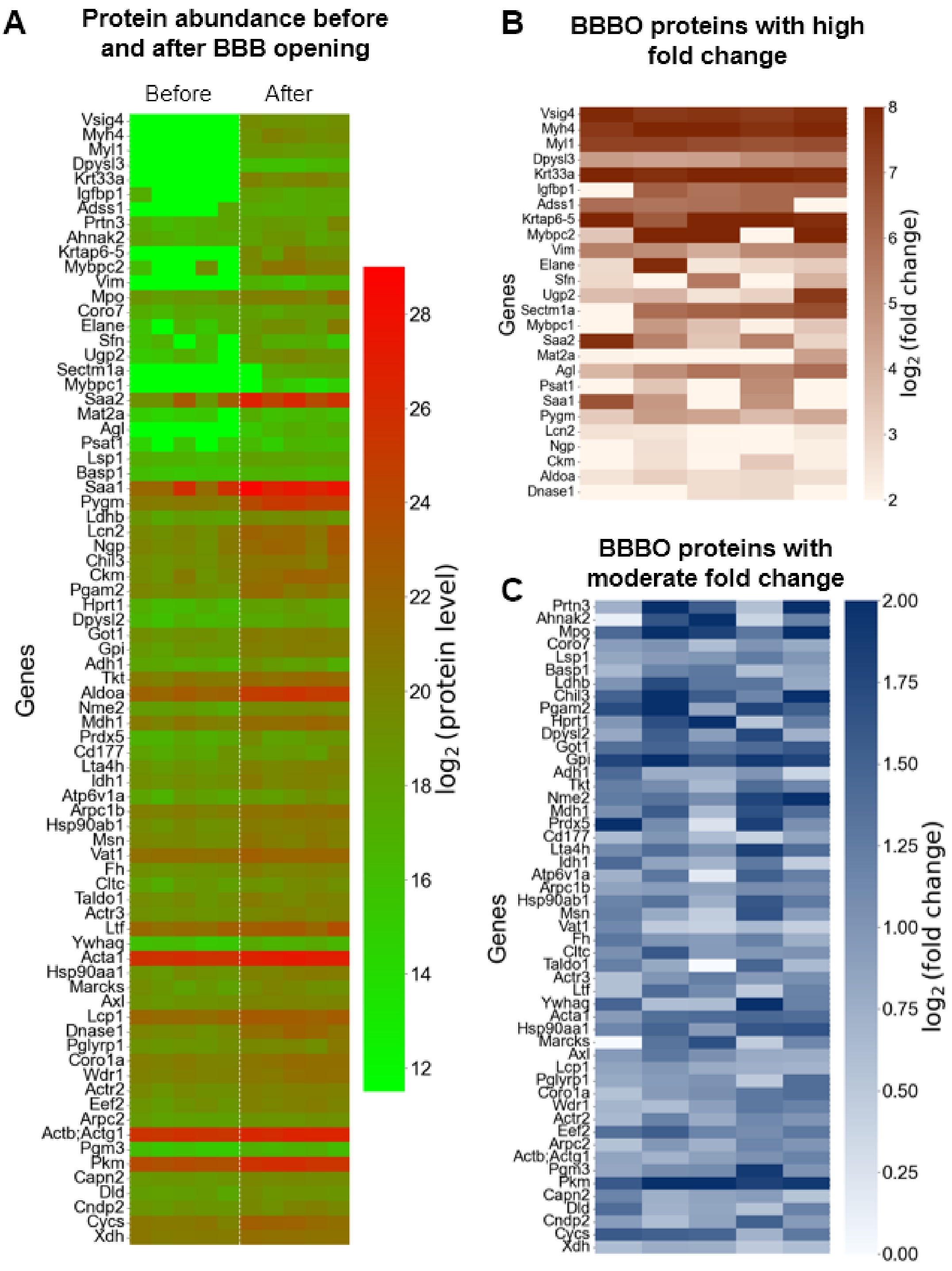


**Supplementary Figure 1. Proteomic markers associated with BBBO in second experiment** (A) Heatmap showing $\log_2$-transformed protein abundance for 77 proteins consistently altered following FUS-mediated BBBO in healthy mice. Differential expression was assessed independently in two experiments using paired *t*-tests comparing pre- and post-BBBO samples. Proteins with a p-value < 0.05 and fold change > 1.5 in both experiments were considered significant. The 77 overlapping proteins are presented here using data from the second experiment, and the corresponding values are provided in supplementary table 2. (B) Subset of BBBO-associated proteins exhibiting large fold changes ($\log_2$ fold change ≥ 2). (C) Subset of BBBO-associated proteins exhibiting moderate fold changes ($\log_2$ fold change < 2).

| Genes | PBS 1 | PBS 2 | PBS 3 | PBS 4 | PBS 5 |
|---|---|---|---|---|---|
| Vsig4 | 8.17 | 7.16 | 7.96 | 7.43 | 6.57 |
| Myh4 | 7.59 | 7.8 | 7.87 | 8.53 | 2.31 |
| Myl1 | 5.28 | 6.93 | 6.97 | 6.43 | 8.03 |
| Dpysl3 | 6.59 | 6.45 | 6.51 | 6.62 | 5.84 |
| Krt33a | 7.33 | 7.44 | 9.21 | 0 | 7.84 |
| Igfbp1 | 7.93 | 5.37 | 6.3 | 6.19 | 5.06 |
| Adss1 | 6.3 | 8.49 | 6.6 | 7.57 | 1.24 |
| Prtn3 | 6.7 | 6.15 | 6.92 | 1.83 | 7.19 |
| Ahnak2 | 6.74 | 5.89 | 6.98 | 1.53 | 6.3 |
| Krtap6-5 | 5.85 | 7.03 | 8.2 | 0 | 5.59 |
| Mybpc2 | 8.05 | 8.39 | 8.23 | 1.17 | 0.63 |
| Vim | 5.88 | 2.41 | 6.29 | 5.74 | 5.97 |
| Mpo | 1.02 | 7.18 | 8.15 | 1.9 | 7.89 |
| Coro7 | 6.16 | 6.9 | 5.39 | 5.54 | 1.88 |
| Elane | 7.28 | 7.16 | 7.66 | 1.66 | 1.83 |
| Sfn | 5.85 | 2.09 | 5.87 | 5.79 | 5.35 |
| Ugp2 | -1.2 | 6.77 | 5.13 | 6.54 | 6.95 |
| Sectm1a | 5.93 | 5.61 | 0.61 | 6.03 | 5.75 |
| Mybpc1 | 5.94 | 5.83 | 3.74 | 0 | 5.9 |
| Saa2 | 3.33 | 2.77 | 4.09 | 4.2 | 7.01 |
| Mat2a | 5.53 | 6.04 | 4.23 | 4.94 | 0.13 |
| Agl | 4.74 | 6.5 | -0.53 | 4.7 | 1.97 |
| Psat1 | 5.4 | 6.09 | 4.66 | 0.29 | 0.67 |
| Lsp1 | 6.67 | 3.33 | 1.73 | 1.62 | 2.74 |
| Basp1 | 4.25 | 4.61 | 4.17 | 1.19 | 0.49 |
| Saa1 | 1.56 | 2.16 | 2.84 | 3.2 | 4.62 |
| Pygm | 2.47 | 3.01 | 1.61 | 2.78 | 2.57 |
| Ldhb | 3.49 | 1.47 | 1.7 | 2.1 | 3.25 |
| Lcn2 | 1.52 | 2.65 | 3.12 | 1.44 | 3.22 |
| Ngp | 2.03 | 2.83 | 3.01 | 1.19 | 2.78 |
| Chil3 | 0.99 | 2.54 | 2.87 | 1.78 | 2.47 |
| Ckm | 1.75 | 3.56 | 1.15 | 2.36 | 1.39 |
| Pgam2 | 0.55 | 2.52 | 1.92 | 2.68 | 1.24 |
| Hprt1 | 2.87 | 2.45 | 1.02 | 0.89 | 1.66 |
| Dpysl2 | 3.41 | 2.58 | 1.37 | 0.53 | 0.95 |
| Got1 | 2.12 | 1.82 | 1.58 | 1.53 | 1.55 |
| Gpi | 1.68 | 3 | 1.64 | 0.61 | 1.37 |
| Adh1 | 3.28 | 1.02 | 1.53 | 1.04 | 1 |
| Tkt | 1.44 | 2.88 | 1.54 | 0.51 | 1.36 |
| Aldoa | 1.18 | 2.85 | 1.11 | 0.84 | 1.32 |
| Nme2 | 1.45 | 2.52 | 1.4 | 0.87 | 0.93 |
| Mdh1 | 0.97 | 2.27 | 1.86 | 1.06 | 1 |
| Prdx5 | 0.38 | 2.61 | 1.32 | 0.84 | 1.96 |
| Cd177 | 0.95 | 1.28 | 1.71 | 1.18 | 1.57 |
| Lta4h | 1.08 | 2.68 | 1.49 | 0.34 | 0.94 |

| | | | | | |
|---|---|---|---|---|---|
| Idh1 | 1.49 | 1.6 | 1.07 | 1.02 | 1.22 |
| Atp6v1a | 2.54 | 1.87 | 1.04 | 0.32 | 0.6 |
| Arpc1b | 1.97 | 1.48 | 0.87 | 0.86 | 1.18 |
| Hsp90ab1 | 1.14 | 2.2 | 1.16 | 0.59 | 1.27 |
| Msn | 0.95 | 2.27 | 1.53 | 0.81 | 0.77 |
| Vat1 | 1.72 | 1.17 | 0.95 | 0.78 | 1.65 |
| Fh | 1.46 | 2.04 | 0.99 | 0.87 | 0.79 |
| Cltc | 1.45 | 2.42 | 1.59 | 0.08 | 0.57 |
| Taldo1 | 0.53 | 1.68 | 1.66 | 1.45 | 0.77 |
| Actr3 | 0.68 | 1.98 | 1.55 | 0.27 | 1.52 |
| Ltf | 0.32 | 1.11 | 2.05 | 0.87 | 1.58 |
| Ywhaq | 0.85 | 2.36 | 1.15 | 0.69 | 0.84 |
| Acta1 | 1.3 | 2.31 | 0.8 | 0.09 | 1.34 |
| Hsp90aa1 | 0.96 | 2.48 | 1.28 | 0.32 | 0.76 |
| Marcks | 1.17 | 1.38 | 1.27 | 1.19 | 0.75 |
| Axl | 1.18 | 0.85 | 1 | 1.49 | 1.09 |
| Lcp1 | 0.28 | 1.28 | 1.28 | 1.4 | 1.31 |
| Dnase1 | 1.29 | 0.4 | 1.54 | 0.94 | 1.36 |
| Pglyrp1 | 0.64 | 0.73 | 1.85 | 0.41 | 1.64 |
| Coro1a | 0.91 | 2.05 | 0.94 | 0.01 | 1.38 |
| Wdr1 | 1.24 | 1.95 | 0.77 | 0.3 | 0.98 |
| Actr2 | 0.91 | 2.04 | 0.84 | 0.16 | 1.24 |
| Eef2 | 0.95 | 1.8 | 1.2 | 0.38 | 0.84 |
| Arpc2 | 0.22 | 1.74 | 1.09 | 0.5 | 1.16 |
| Actb;Actg1 | 1.01 | 1.84 | 0.52 | -0.04 | 1.36 |
| Pgm3 | 1.88 | 1.47 | 0.23 | 0.71 | 0.39 |
| Pkm | 0.47 | 1.6 | 0.73 | 0.54 | 1.14 |
| Capn2 | 1.3 | 1.23 | 0.73 | 0.33 | 0.8 |
| Dld | 0.78 | 0.61 | 0.88 | 0.82 | 1.2 |
| Cndp2 | 0.35 | 1.2 | 1.68 | 0.88 | 0.11 |
| Cycs | 0.03 | 0.54 | 0.97 | 1.26 | 1.08 |
| Xdh | 0.45 | 1.15 | 0.2 | 0.36 | 1.09 |

**Supplementary Table 1.** Differential expression of proteins identified in the first sonobiopsy experiment

| Genes | PBS 1 | PBS 2 | PBS 3 | PBS 4 | PBS 5 |
|---|---|---|---|---|---|
| Vsig4 | 7.95 | 7.48 | 7.62 | 7.4 | 7.75 |
| Myh4 | 7.5 | 8.63 | 8.25 | 7.68 | 7.96 |
| Myl1 | 7.2 | 7.18 | 6.91 | 6.73 | 6.92 |
| Dpysl3 | 4.58 | 4.38 | 4.46 | 5.05 | 5.34 |
| Krt33a | 8.56 | 7.73 | 8.31 | 8.63 | 7.84 |
| Igfbp1 | 1.06 | 6.33 | 5.82 | 6.13 | 6.19 |
| Adss1 | 5.94 | 5.73 | 5.87 | 6.13 | -0.24 |
| Prtn3 | 0.74 | 2.32 | 1.55 | 0.58 | 2.05 |
| Ahnak2 | 0.12 | 1.64 | 2.32 | 0.38 | 1.19 |
| Krtap6-5 | 8.18 | 6.5 | 9.02 | 7.99 | 7.78 |
| Mybpc2 | 3.39 | 9.47 | 9.74 | 1.01 | 8.82 |
| Vim | 5.41 | 5.01 | 4.12 | 5.14 | 5.22 |
| Mpo | 1.44 | 2.35 | 1.81 | 1.3 | 2.12 |
| Coro7 | 0.93 | 1.04 | 0.61 | 1.02 | 0.8 |
| Elane | 2.82 | 7.86 | 2.45 | 2.86 | 3.23 |
| Sfn | 2.86 | 0.48 | 5.57 | 1.19 | 3.93 |
| Ugp2 | 3.66 | 3.88 | 2.54 | 3.07 | 7.48 |
| Sectm1a | 0 | 5.92 | 6.28 | 6.5 | 6.83 |
| Mybpc1 | 0 | 4.68 | 3.53 | 2.2 | 3.39 |
| Saa2 | 7.71 | 5.36 | 3.43 | 5.34 | 2.98 |
| Mat2a | 2.09 | 1.44 | 1.1 | 1.33 | 4.48 |
| Agl | 3.83 | 5 | 5.76 | 5.28 | 5.98 |
| Psat1 | 1.66 | 3.43 | 0.96 | 5.06 | 1.35 |
| Lsp1 | 0.83 | 0.94 | 0.98 | 1.26 | 0.99 |
| Basp1 | 0.65 | 1.18 | 1.29 | 0.58 | 0.85 |
| Saa1 | 6.7 | 4.7 | 1.52 | 4.89 | 1.89 |
| Pygm | 3.26 | 4.56 | 4.35 | 3.7 | 4.21 |
| Ldhb | 1.01 | 1.72 | 1.3 | 1.31 | 0.8 |
| Lcn2 | 2.58 | 2.47 | 1.99 | 1.73 | 2.46 |
| Ngp | 1.59 | 2.66 | 2.04 | 1.73 | 2.19 |
| Chil3 | 1.5 | 2.06 | 1.55 | 1.16 | 1.99 |
| Ckm | 1.72 | 2.63 | 1.51 | 3.28 | 2.21 |
| Pgam2 | 1.72 | 2.28 | 0.83 | 1.78 | 1.53 |
| Hprt1 | 0.99 | 1.7 | 2.11 | 0.53 | 1.24 |
| Dpysl2 | 0.82 | 1.54 | 0.92 | 1.75 | 0.73 |
| Got1 | 1.37 | 1.49 | 1.3 | 1.43 | 1.59 |
| Gpi | 1.78 | 2.11 | 1.59 | 1.9 | 1.79 |
| Adh1 | 1.42 | 0.76 | 0.76 | 1 | 0.37 |
| Tkt | 1.22 | 1.08 | 0.65 | 1.4 | 1.15 |
| Aldoa | 2.52 | 3.08 | 2.73 | 2.82 | 2.69 |
| Nme2 | 1.25 | 1.34 | 1.08 | 1.79 | 1.99 |
| Mdh1 | 1.05 | 1.56 | 0.65 | 1.67 | 1.41 |
| Prdx5 | 2.61 | 1.09 | 0.24 | 1.84 | 1.05 |
| Cd177 | 0.64 | 1 | 0.61 | 0.4 | 0.86 |
| Lta4h | 1.11 | 1.39 | 1.03 | 1.83 | 1.4 |

| | | | | | |
|---|---|---|---|---|---|
| Idh1 | 1.43 | 0.86 | 0.71 | 1.28 | 0.46 |
| Atp6v1a | 0.74 | 1.3 | 0.18 | 1.53 | 1.22 |
| Arpc1b | 0.87 | 0.93 | 0.88 | 1.08 | 1.05 |
| Hsp90ab1 | 1.18 | 1.34 | 0.75 | 1.61 | 1.27 |
| Msn | 1.22 | 0.79 | 0.46 | 1.65 | 0.94 |
| Vat1 | 1.13 | 0.48 | 0.42 | 0.65 | 0.43 |
| Fh | 1.3 | 1 | 0.93 | 1.2 | 0.75 |
| Cltc | 1.05 | 1.6 | 0.95 | 0.99 | 1.03 |
| Taldo1 | 1.22 | 0.81 | -0.02 | 1.47 | 0.65 |
| Actr3 | 0.58 | 0.99 | 1.23 | 0.97 | 1.06 |
| Ltf | 0.58 | 1.41 | 1.12 | 0.48 | 1.19 |
| Ywhaq | 1.48 | 0.6 | 0.62 | 2.08 | 1.2 |
| Acta1 | 0.94 | 1.37 | 1.41 | 1.46 | 1.4 |
| Hsp90aa1 | 1.15 | 1.48 | 0.94 | 1.62 | 1.65 |
| Marcks | -0.09 | 1.33 | 1.68 | 0.46 | 1.16 |
| Axl | 0.95 | 1.28 | 1.05 | 0.78 | 0.77 |
| Lcp1 | 0.85 | 0.98 | 0.78 | 0.74 | 0.71 |
| Dnase1 | 1.53 | 1.83 | 2.71 | 2.84 | 1.55 |
| Pglyrp1 | 0.8 | 0.94 | 1.03 | 0.48 | 1.39 |
| Coro1a | 0.56 | 0.94 | 1.07 | 1.29 | 1.38 |
| Wdr1 | 0.69 | 0.61 | 0.89 | 1.29 | 1.2 |
| Actr2 | 0.65 | 1.19 | 0.78 | 1.13 | 1 |
| Eef2 | 1.37 | 1.55 | 1.17 | 1.08 | 1.28 |
| Arpc2 | 0.56 | 0.98 | 0.74 | 1.16 | 1 |
| Actb;Actg1 | 0.85 | 0.71 | 0.89 | 1.03 | 1.18 |
| Pgm3 | 0.83 | 1.07 | 1.1 | 1.89 | 1.01 |
| Pkm | 1.6 | 2.42 | 2.14 | 1.82 | 1.91 |
| Capn2 | 1.17 | 0.75 | 0.87 | 0.93 | 0.54 |
| Dld | 1.41 | 0.73 | 0.84 | 0.55 | 1.2 |
| Cndp2 | 0.92 | 0.59 | 0.86 | 1.52 | 0.95 |
| Cycs | 1.58 | 1.51 | 1.47 | 0.72 | 1.28 |
| Xdh | 0.62 | 0.78 | 0.76 | 0.48 | 0.61 |

**Supplementary Table 2.** Differential expression of proteins identified in the second sonobiopsy experiment
